# Advances and perspectives in tissue clearing using CLARITY

**DOI:** 10.1101/144378

**Authors:** Kristian H. Reveles Jensen, Rune W. Berg

## Abstract

CLARITY is a tissue clearing method, which enables immunostaining and imaging of large volumes for 3Dreconstruction. The method was initially time-consuming, expensive and relied on electrophoresis to remove lipids to make the tissue transparent. Since then several improvements and simplifications have emerged, such as passive clearing (PACT) and methods to improve tissue staining. Here, we review advances and compare current applications with the aim of highlighting needed improvements as well as aiding selection of the specific protocol for use in future investigations.

## Chemical compounds

Chemical compounds mentioned in this article: 1-Ethyl-3-(3-dimethylaminopropyl)carbodiimide (PubChem CID: 15908); 2,2’-Thiodiethanol (PubChem CID: 5447); *α*-Thioglycerol (PubChem CID: 7291); Acrylamide (PubChem CID: 6579); D-Sorbitol (PubChem CID: 5780); Diatrizoic acid (PubChem CID: 2140); Gylcerol (PubChem CID: 753); Iohexol (PubChem CID: 3730); Iomeprol-d3 (PubChem CID: 46781978); N,N,N’,N’-Tetrakis(2-Hydroxypropyl)ethylenediamine (amino alcohol, PubChem CID: 7615)

## Introduction

The process of clearing tissue for the purpose of histological analysis has recently become a common tool in biological investigations. The purpose is to keep the proteins in a structure while removing the light-scattering lipids, and thus to allow deep penetration of light for fluorescent 3D microscopy. Several clearing methods have recently been developed.

Here, we focus on the advancement in the procedure called CLARITY (*C*lear *L*ipid-exchanged *A*crylamidehybridized *R*igid *Imaging/Immunostaining/I*n situ-hybridization-compatible *T issue-hY drogel)*. It is a convenient histological fixation and clearing technique that enables immunohistochemistry (ICH) and maintenance of fluorophores during imaging of large volumes for 3D-reconstruction. The method was developed by Chung, Gradinaru, Deisseroth & colleagues and relies on the removal of lipids while keeping the protein and DNA of the tissue by creating a hydrogel by cross-linking with acrylamide (Chung *et al.*, 2013; Chung and Deisseroth, 2013). Since then several improvements and simplifications have emerged, such as passive clearing (PACT), and other adaptations (EDC-CLARITY).

We review advances, and compare current applications and limitations for this methodology, with the aim of highlighting needed improvements as well as aiding selection of the specific CLARITY or hydrogel-based protocol for use in future investigations. CLARITY was originally demonstrated on rodent brains, but has since been successfully applied to other tissues organs and species, e.g. fish and plants. In this review, we primarily focus on nervous tissue although the protocols work for other tissues as well.

## The original CLARITY method

CLARITY works by polymerising proteins, DNA, and RNA in fixed tissue using acrylamide to form a tissue-hydrogel hybrid before lipid-removal. The acrylamide based hydrogel binds molecules with amine ends, predominantly proteins and a small proportion of DNA and RNA, into a skeleton structure, which is permeable to larger molecules. The lipids are then encapsulated into micelles using detergent (sodium dodecyl sulfate, SDS). The micelles are electrically charged and can therefore easily be carried out through the pores by an applied electrical field. Antibodies for immunohistochemistry (ICH) can also penetrate the hydrogel to reach and stain the fixed membrane proteins. The original protocol was first demonstrated on whole adult mice brains and can be summarised in the following steps:

1. 1. **Fixation of tissue.** The animal is transcardially perfused with a mixture of 4% (w/v) paraformaldehyde (PFA), 4% (w/v) acrylamide, 0.05% (w/v) bis-acrylamide, 0.25% (w/v) VA-044 and in phosphatebuffered saline (PBS), and the brain is removed and incubated in the same solution for 3 days.
2. 2. **Hydrogel formation.** A 3-hour heat activation at 37°C for the thermal initiator, VA-044, results in the polymerisation of the acrylamide and bis-acrylamide to the PFA-fixed tissue.
3. 3. **Lipid extraction.** Tissue clearing, or lipid removal, is accomplished by incubating the tissue in a sodium borate buffer (0.2 M, pH 8.5) containing 4% (w/v) SDS in a custom-designed electrophoretic chamber. The basic pH leads to negatively charged micelles, which pass trough the porous hydrogel by electrophoresis using a 10-60 *V* current applied across the sample at 37-50°C for two days.
4. 4. **Refractive index matching.** The tissue is washed in PBS for two days to remove SDS. The tissue-hydrogel has a refractive index (RI) of 1.44-1.46. The tissue is incubated with a proprietary imaging solution, FocusClear (CelExplorer Labs Co.), with a similar RI of 1.454 to increase transparency.

The novelty in the CLARITY method is the second step since utilization of a hydrogel had a clear benefit compared with other methods. Even using the harsh 4% SDS for a week, the cleared tissue had only 8% protein loss. For comparison, in PFA-fixed tissue in 4% SDS, without the hydrogel up to 65% protein was lost. Similar protein loss is observed using other clearing methods without a hydrogel, such as Sca/e, which uses urea and glycerol (Hama *et al.*, 2011). The effective retention of proteins and antigens also enabled repetitive staining, where antibodies could be washed out briefly in 4% SDS and restained.

However, the step of lipid extraction had several limitations: active clearing with electrophoresis required a custom-designed chamber with a continuous exchange of SDS solution. The electrical current, if not properly controlled, can decompose or discolour the tissue. Thus, since the original CLARITY method both required expensive elements in step 4 such as the RIMS medium (FocusClear) and was difficult to implement, a potential for improvement presented itself. Several labs have employed a more simple version of CLARITY without electrophoresis, which is the passive CLARITY clearing. The need for cheaper refractive index imaging solutions than the original (FocusClear) has also become apparent. First, we will address the general challenges in tissue clearing.

“

CLARITY in its original form used electrophoretic tissue clearing (ETC) to extract lipids from large samples, which can be challenging to implement and can cause variability in final tissue quality, including epitope loss, damage to fine processes, and tissue browning due to heating (Lee *et al.*, 2014).

### General challenges

There are several issues to consider when clearing tissue. Processing time, the number of procedural steps, the toxicity of reagents, signal and bleaching of fluorophores and the required concentration of primary and secondary antibodies, which are usually expensive and therefore beneficial to reduce. Efficiency in the extraction of lipids from the hydrogel, i.e. how much of the original lipids remain and how much time does the procedure require. Successful extraction of lipids is usually manifested in the transparency of the sample and the incubation time is often adjusted by the investigator for the sample to achieve the appropriate properties. Although the tissue transparency has been attributed to the removal of lipids during the procedure, it is still unknown to what extent the lipids are cleared from the sample and how much remains in transparent samples. This is contrary to the degree of protein loss, which has been quantified (Chung *et al.*, 2013; Lai *et al.*, 2016).

A major challenge introduced by clarified tissue samples is the imaging technology followed by the data analysis. Using conventional single photon laser scanning confocal microscopy, vast areas of the sample become unnecessarily illuminated, which introduces photobleaching of the fluorophores as a general issue. Photo-bleaching is potentially devastating in samples with a weak fluorescent signal, such as in Brainbow tissue (Cai *et al.*, 2013).

For the same reason, it is important to limit the protein loss during storage of CLARITY samples. A novel imaging technology overcomes this issue by selectively only illuminating a single thin plane in a sequential manner. This technique is known as light-sheet microscopy, which can give a high rate of data acquisition of high contrast and resolution, and with very limited exposure to the rest of the sample (Stefaniuk *et al.*, 2016). Light-sheet microscopy is remarkably faster than traditional single-or two-photon laser scanning microscopy. Several variants of light-sheet microscopy are commercially available, but they are not inexpensive and will represent a major investment to most laboratories. Another issue to consider is the price of the equipment and protocol compounds. The equipment necessary to set up active CLARITY with electrophoresis is approximately 6,800 USD. FocusClear is the originally recommended imaging media, which is expensive, but other less expensive alternative exist such as RIMS, sRIMS and glycerol.

It is also important to consider that acrylamide, the primary component in the formation of the hydrogel, is a carcinogenic compound and is on the U.S. federally regulated list of ‘extremely hazardous substances’. Finding a less toxic alternative is therefore appealing. An acrylamide-free procedure with fewer steps has been suggested that the SDS alone can extract the lipids without the need for the toxic hydrogel (Lai *et al.*, 2016).

Based on these issues, we have composed a list of improvements in the CLARITY protocol below.

## Variants and improvements of CLARITY

The steps of the CLARITY procedure including staining and imaging is summarized as follows: 1) Tissue fixation and cutting, 2) Hydrogel polymerization, 3) Passive or active lipid removal, 4) Staining, 5) Optical clearing, 6) Imaging (figure 1). Here we summarize the improvements in these steps. A list of variants of the CLARITY improvements are found in table 3.

**Figure 1:**
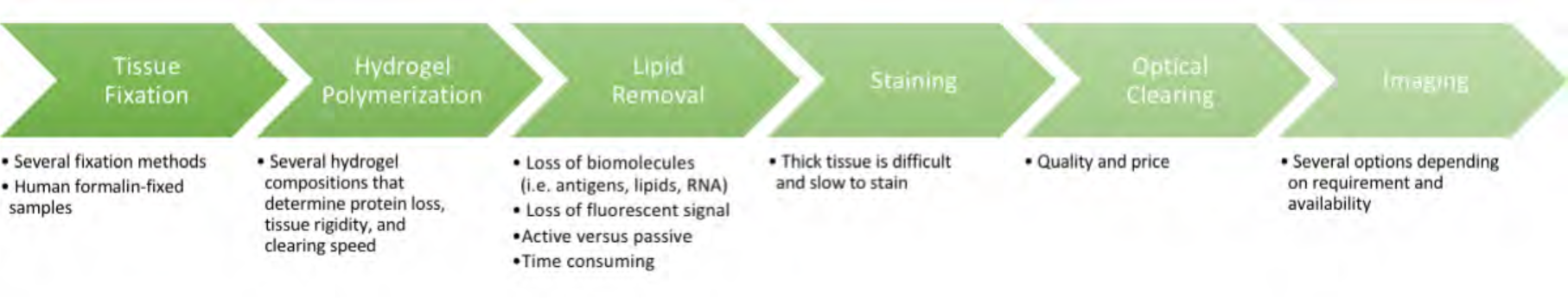
Steps in the CLARITY procedure including challenges and considerations.

### 1. Tissue fixation and cutting

Tissue fixation is the initial step in practically all histology as well as CLARITY. The original protocol prescribes the animal to be transcardially perfused with 4% PFA together with the acrylamide monomers and activator VA-044, and the nervous tissue further incubated in the same solution for 3 days (Chung *et al.*, 2013). In the following published protocol, the incubation was shortened to 1 day (Tomer *et al.*, 2014). However, fixation with 4% PFA alone was later demonstrated as sufficient for the subsequent cross-linking and hydrogel polymerization if the tissue was just incubated in the monomer solution afterwards (Costantini *et al.*, 2015; Lee *et al.*, 2014; Zheng and Rinaman, 2016; Jensen and Berg, 2016). Transcardial perfusion of PBS to remove blood followed by the fixative or monomer solution is preferable. However, for some purposes just fixing the excised tissue, which has not been perfused, e.g. human tissue, overnight in 4% PFA in PBS is sufficient. Trimming the tissue down to the minimal required volume speeds up clearing and staining. Smaller tissue volume also results in less light scattering and clearer imaging. This can be done by embedding the tissue in low melting agar and cutting the tissue down to the most practical form e.g. a hemisphere, 1-2 mm slices or sections on a vibratome. Slicing the tissue after the hydrogel formation (step 2) is not recommended since the hydrogel swells to a sticky mass and is difficult to cut accurately, besides being carcinogenic. We recommend cutting before incubation and the polymerisation step.

### 2. Hydrogel polymerization and composition

Warning! Acrylamide is carcinogenic and should be handled with great care. After fixation, the tissue needs to be cross-linked and hybridised into a hydrogel by the acrylamide monomers to stabilise biomolecules and retain them during tissue clearing. The purpose is to achieve a fast and homogeneous clearing process, with good antibody penetration depth and retaining the desired antigens (protein). PFA fixes the tissue and supposedly acts as an anchor point for the acrylamide monomers to polymerise into a mesh. After removal of lipids the proteins are left in a hydrogel (step 3).

The original hydrogel monomer solution (4% PFA, 4% acrylamide, 0.05% bis-acrylamide) was optimized to provide a balance between hydrogel rigidity and porosity with minimal protein loss (∼8% when cleared in a 4% SDS borate buffer) after ETC clearing (Chung *et al.*, 2013). However, solution adjustments may be useful for applying CLARITY to certain kinds of tissues and passive and active clearing.

While an emphasis has been put on optimising the hydrogel composition for passive clearing, the modifications in the hydrogel composition are applicable to both passive and active clearing by electrophoresis. The hydrogel presented in the PACT method (the A4P0 composition) is the current default in for passive clearing, but it is also applied in active clearing with the ACT-PRESTO protocol. Others have used 2%PFA and 2% acrylamide for active clearing (Bastrup and Larsen, 2017). The variations in the hydrogel monomer solution are therefore presented here before addressing the lipid extraction step.

### PACT and PARS

Yang *et al.* (2014) focused on developing a simple version of CLARITY based purely on the passive removal of lipids, i.e. passive clearing. They tested several hydrogel compositions and SDS concentrations leading to an improved protocol named PACT (*pa*ssive *C*LARITY *t*echnique) protocol. They had three main findings: First, omitting PFA in the monomer solution did not change the degree of protein loss, but did increase IgG penetration depth i.e. better antibody labelling. Second, omitting PFA leads to faster transparency, but also a greater tissue swelling (∼230%), yet the swelling is transient, and may improve staining by making the tissue more porous. Finally, lower acrylamide concentration (i.e. 2%) leads to less cross-linking and less solid tissue, this in turns results in faster clearing, but also greater tissue swelling and protein loss. Hence, they concluded that 4% acrylamide, which they referred to as A4P0 solution, is optimal for nervous tissue and other tissues e.g. liver and kidney.

While A4P0 was optimised for passive clearing, the recipe is also used in active clearing (Lee *et al.*, 2016). Rather than tissue incubation in the monomer solutions, the PACT reagents can be delivered either via the vasculature (intracardial injection) or intracranially by injection into the cisterna magna or a subdural cannula above the olfactory bulb to achieve whole body or CNS clearing and labelling. The technique is termed PARS (*P*erfusion-assisted *A*gent *R*elease in *S*itu). The PACT reagents used in PARS are recirculated into the cerebrospinal fluid (CSF) or through the whole-body vasculature for several days-to-weeks in closed-loop perfusion system mimicking regular blood flow allowing for clearing and staining deep in the tissue (Tomer *et al.*, 2014). PARS does not change the speed of tissue clearing of the brain or internal organs (Woo *et al.*, 2016), table 2. The PARS delivery system can be combined with other of the below methods if needed. In summary, PACT/PARS is a radical improvement since they provide a simpler recipe, which also gives a faster clearing with better antibody staining.

### psPACT

psPACT (*processes separate* PACT) enhanced the clearing speed by splitting the monomer incubation into *separate processes* before polymerisation: an initial 24 hr incubation at 37^o^C in acrylamide solution (4% w/v in PBS), followed by a 6-12 hr incubation at room temperature in VA-044 initiator solution (0.25% w/v in PBS) for the hydrogel formation. This protocol shortens the subsequent clearing time by ∼10% (Woo *et al.*, 2016). A comparison of protein loss with the PACT method has not yet been performed.

### ePACT

ePACT (*expansion-enhanced* PACT) is used to expand thin tissue sections (100µm) to larger volumes for better resolution and resolution beyond the diffraction limit (Treweek *et al.*, 2015). The hydrogel is composed of an acrylate-acrylamide copolymer and cleared with 10% SDS, where IHC such be done prior, and then digested with collagenase before incubation in water, which results in water absorption and tissue expansion.

### FASTClear

In CLARITY protocols proteins are assumed to be fixed to an acrylamide scaffold via PFA, whereas lipids are washed out by the amphiphilic SDS-micelles. Nevertheless, Lai *et al.* (2016) demonstrated an acrylamide-free version of CLARITY, where the tissue is essentially only washing with SDS followed by RIMS, gave better results for formalin-fixed tissue. Therefore Lai *et al.* (2016) suggested that the tissue–PFA–acrylamide cross-linking does not actually occur, and therefore omitting acrylamide-polymerisation is not only easier, but also better. Hence, they developed a Free of acrylamide SDS-based tissue-clearing (FASTClear) protocol (Liu *et al.*, 2016).

### Other modifications

It is worth noting that the PACT protocol also omits bis-acrylamide, which acts as a secondary cross-linker, and therefore does not link the PFA-tagged biomolecules, but rather directly cross-links poly-acrylamide chains to form the gel. Therefore Bis-acrylamide increases the rigidity of the hydrogel network by creating cross-links inside cavities that may be void or sparse of biomolecules such as the ventricles. Bis-acrylamide also causes all the hydrogel solution surrounding the tissue sample to cross-link and form a gel during the polymerization step, which is a helpful aid in determining successful polymerisation. The surrounding gel can easily be removed manually from the tissue by physical rubbing/handling. Without bis-acrylamide and the gel formation outside the tissue, the sample can simply be removed from the solution following the polymerization step and is therefore recommended for small or fragile specimens that cannot withstand the physical gel removal process.

The incubation time of the tissue in monomer solution before polymerisation varies among authors with 1-3 days of incubation for a whole rodent brain, while for 1-2 mm slices overnight is sufficient. Saponin is a mild non-ionic surfactant often used to permeabilise cellular membranes in conventional histology. It was briefly mentioned by Chung and Deisseroth (2013) as an adjunct in the monomer solution to improve diffusion of the hydrogel monomer and initiator into tissues, where cardiac perfusion is not possible e.g. post-mortem human tissues and zebrafish brains. Saponin reportedly shortens the incubation time required in the hydrogel monomer infusion process. However, saponin may cause bubbles, and therefore it is not recommended and red has only been reported utilised in a study of ovarian follicles (Feng *et al.*, 2017). In zebrafish adding dimethyl sulfoxide (DMSO) improves monomer penetration.

Another challenge for users has been initiating the polymerisation, which is inhibited by oxygen. The original CLARITY protocol, therefore, involved a vacuum desiccation chamber(Chung *et al.*, 2013). While Yang *et al. (2014)* demonstrated it was sufficient to displace the oxygen from the solution by bubbling for ∼10 min with Nitrogen gas before initiation. Some users have found it sufficient to prevent aeration by sealing the monomer solution off by placing a layer of vegetable oil on top in the test tube (forum.claritytechniques.org), yet others simply filled the tube entirely with the monomer solution to minimise inhibitory effects of the oxygen (Bastrup and Larsen, 2017). Another method simply uses double the concentration of the initiator, VA-044, which eliminates any vacuum purging or nitrogen backfilling. This method is called ‘simplified CLARITY method’ (or SCM), but was only tested on thin sections, of 100 µm(Sung *et al.*, 2016).

## 3. Lipid removal

Since membrane lipids are the main cause of light diffraction in tissues and comprise ∼60% of nervous tissue, lipid elution renders the tissue translucent. Temperature and solutions affect lipid clearing: speed, uniformity and protein loss. The original CLARITY protocol used a 4% SDS (sodium borate buffer, pH 8.5) intended for active clearing by electrophoresis (see below). Tissue clearing can be *passive* where lipids captured in detergent micelles slowly diffuse out of the tissue into the wash solution or *active* at an accelerated rate by applying an electric field (figure 2). In a study comparing active and passive CLARITY, there were no significant differences in protein concentration as an effect of active versus passive clearing (Epp *et al.*, 2015).

**Figure 2:**
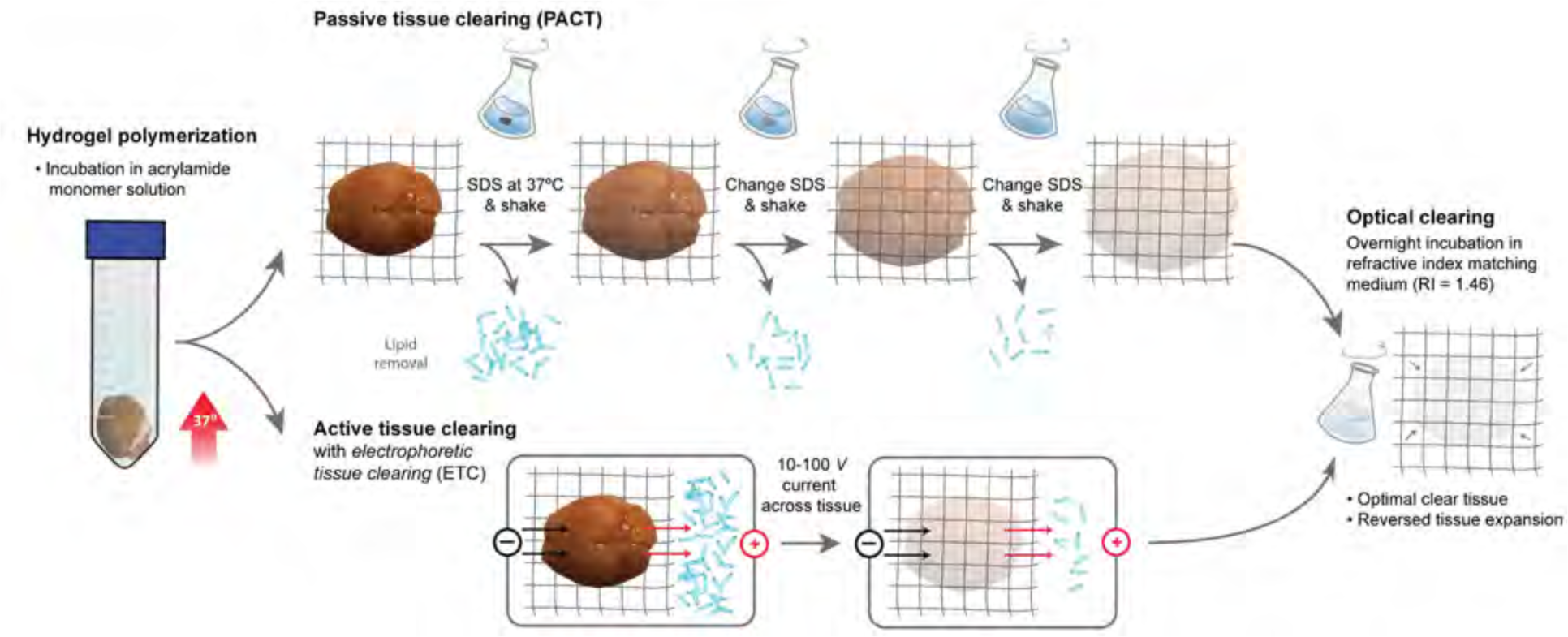
CLARITY tissue clearing: passive and active. The hydrogel-tissue hybrid is formed (left) by an incubation in the acrylamide monomer solution with the initiator VA-044. Polymerization is activated by heat. Afterwards, the lipids can either be cleared from the tissue by passive (top) or active (bottom) methods. The detergent, SDS, collects lipids and carries them out of the tissue leaving behind a transparent hydrogel-tissue hybrid. Passive tissue clearing, PACT (top), is straightforward and inexpensive, but also a slower process as it relies on diffusion, which is enhanced by gentle shaking and an elevation of temperature. During this process the tissue absorbs water and expands in volume up to ∼200%, which improves the access of antibodies and thus the staining. The swelling is reversed in the hygroscopic medium (right). The alternative method of tissue clearing requires more equipment and supervision as it uses electrophoresis (bottom). The negatively charged lipid micelles are actively carried by the electrical field, which greatly enhance the extraction rate. In the final step of optically clearing the tissue (right) the hydrogel is incubated overnight in a refractive index matching medium to complete transparency.

### Passive lipid removal

Passive clearing without electrophoresis is gentle on the tissue with little risk of damage. The clearing lasts longer, which can reduce the signal from fluorescent proteins and increase the loss of biomolecules. It is inexpensive and requires little equipment. A simplified diffusion-based method omitting the electrophoretic chamber, coined CLARITY2, to clarify <1.5 mm thick slices was proposed by Poguzhelskaya *et al.* (2014).

### PACT

Yang *et al.* (2014) optimised the hydrogel composition for passive clearing and also tested several SDS concentrations leading to the improved PACT protocol. They found that 8% SDS (in PBS buffer, pH 7.5) gave a faster and more uniform tissue clearing compared to 4% SDS, 20% SDS and 10% sodium deoxycholate. For this reason, PACT has become the default passive CLARITY protocol. There is no significant difference in the swelling and shrinking behaviour of passively cleared rodent gut tissue with CLARITY or PACT solution, nor a difference in clearing time (Neckel *et al.*, 2016).

### mPACT

Woo *et al.* (2016) was inspired by the SeeDB method (Ke *et al.*, 2013) and added 0.5% *α*-thioglycerol to the PACT clearing solution, which improved clearing time with ∼10%. They named it psPACT (process–seperated PACT) with added *α*-thioglycerol mPACT (*modified* PACT). The mPACT gives a combined ∼25% faster clearing compared with regular PACT on most tissues (table 2). Theeffect of *α*-thioglycerol on protein loss has not been reported, but any protein loss is unlikely to have an adverse effect. On the contrary, since *α*-thioglycerol preserves proteins by counteracting the Maillard reaction, i.e. browning of the specimen, autofluorescence and antigen loss is likely to be minimal.

### Temperature and speed

Tissue clearing is often performed at room temperature (RT) or 37°C. Generally, increasing temperature accelerates clearing and imaging depths but at the risk of damage to the tissue and quenching fluorescent proteins. For instance, passively clearing of 1 mm cortical sections takes ∼48 hours at 37°C compared to ∼12 hours at 57°C(Yu *et al.*, 2017). Yu *et al.* (2017) found that an elevated temperature range of 42-47°C for PACT resulted in faster transparency and deeper imaging depth than 37°C without affecting the fluorescence signal. There is reportedly no significant difference in protein loss at higher temperatures such as 55°C compared to 37°C(Epp *et al.*, 2015).

Passive clearing can be improved by gentle shaking, continuously replacing the clearing solution (figure 2) or with a flow-assisted clearing setup with a circulator. An alternative setup without a circulator is using a 50 ml conical tube perforated with several holes at the 15-20 ml mark and at the bottom of the tube, which is inserted into a 250 ml glass bottle filled with clearing solution. Unidirectional flow is created by using a magnetic stir bar on a stirring hot plate to accelerate the clearing at the desired temperature (capture-clarity.org/optimized-clarity). The clearing time of various tissues and protocols are listed in table 2.

### Active lipid removal by electrophoresis

Tissue is composed of different types of lipids primarily fatty acids and phospholipids. The detergent, SDS, captures the lipids in SDS micelles. The micelles are negatively charged at basic pH (7.5–8.5) and carries the lipids along the electric field. This electrophoretic tissue-clearing (ETC) enhances the speed of extraction by orders of magnitude such that a large tissue sample e.g. a mouse brain becomes transparent a matter of hours to days instead of days to weeks during passive clearing (Tomer *et al.*, 2014). Lipid clearing using ETC is faster and therefore results in less tissue swelling compared with passive clearing.

The ECT system should have three elements: 1) The chamber containing the sample and electrodes. 2) A circulator, which controls flow rate and temperature. This can be a simple pump and a liquid reservoir that serves to remove electrophoretic by-products such as acid, bubbles and heat. 3) A buffer filter to filter out larger particles in the clearing solution.

There are, however, important caveats to consider when using ETC. If the electrical field is too strong, small bubbles will form inside the hydrogel and make the hydrogel opaque, while bigger bubbles may rupture hydrogel (Kim *et al.*, 2015). Keeping the voltage small enough to avoid this issue, ETC is a helpful step in clearing the sample. The electric field is recommended to be applied as a low–voltage constant current generator of 250-280 mA to ensure low voltage field (10-40 *V*). Such a field will avoid electrolysis of water in the liquid and formation of bubbles (Lee *et al.*, 2014). Running ETC at higher temperatures (55°C) produces very clear tissue, but the tissue tends to lose structural integrity (Epp *et al.*, 2015). Lowering the temperature ∼15°C to counteract Joule–heating is also recommended (Kim *et al.*, 2015). Practically the tissue sample is placed in a custom-designed electrophoretic chamber with platinum electrodes as well as drainage and access of liquid (Tomer *et al.*, 2014; Lee *et al.*, 2016). The original CLARITY chamber had wire electrodes with a surface area of 314mm^2^, which could clear a mouse brain in 5-16 days, whereas electrode plates with a greater surface area of 1200mm^2^as in ACT gives faster clearing (whole mouse brain and other organs within 24 hrs, table 2) and less tissue damage (Lee *et al.*, 2016).

Another caveat of ECT is the flux of lipids is in one direction along the field, which means that the central parts, as well as the parts along borders in parallel with the field, will not have extraction lipids in the same degree as the perpendicular parts. As a result, there is a non-even lipid–extraction and penetration of antibodies when using ECT for ICH. A solution to this problem is, rather than using a static field, to change the direction of the electric field over time, e.g. by rotating the direction. Such rotational electrical field enhances the stochastic electro-diffusion, which enhances lipid extraction as well as staining of large and dense tissue with nuclei dyes, proteins, antibodies (Kim *et al.*, 2015). The two-chamber design with the rotational electrical field can clear mice brains and other organs within 3 days (table 2).

While most chamber designs are freely and commercially available, setting up ETC can be expensive. 3D–printing of shared designs may significantly reduces costs for experimenters and allow for organ-specific designs (Sulkin *et al.*, 2013; Miller and Rothstein, 2017); e.g. a tissue cutting matrix (Tyson *et al.*, 2015) and imaging chambers (idisco.info/idisco-protocol). An overview of the three ECT approaches and where to find free designs and commercial ETC chambers are summarised (table 1).

**Table 1:** Active lipid removal using electrophoretic tissue clearing (ETC): protocols and chambers.

| Protocol | Electrodes and field | Speed and tissue damage | Time to clear a mouse brain | Freely available protocol and chamber blueprints | Commercial options |
| --- | --- | --- | --- | --- | --- |
| CLARITY | Wire-electrodes; unidirectional field (10-30 V; 280mA) | (+) Linear with $V$<br>Discolouration and tissue damage | 15–16 days | Chung *et al.* (2013); Tomer *et al.* (2014)<br>Bastrup and Larsen (2017)<br>wiki.claritytechniques.org | |
| ACT | Large surface plate-electrodes; unidirectional field (1.5A) | (+++ Linear with $V$<br>Less tissue damage | 6 hours | Lee *et al.* (2016) | X-CLARITY (Logos Biosystems) |
| Stochastic Electrotransport* | Wire-electrodes; rotating field (10-100 V) | (++) Quadratic with $V$<br>Least tissue damage | 3 days | Sylwestrak *et al.* (2016) and www.chunglabresources.com | SmartClear I & II (LifeCanvas Tech) |

\*The Stochastic Electrotransport system can also be used to rapidly deliver dyes or antibodies into the tissue during staining.

### Loss of fluorescent proteins and antigens

Many investigations employ viral tracers to induce fluorescent protein expression in specific neuronal populations (figure 3a). However, the fluorescent proteins are vulnerable to over-fixation, denaturation and elution with subsequent signal loss and the fluorescence may require amplification. Another common tool is to image fluorescent reporters such as green–or yellow fluorescent proteins (GFP or YFP), which are expressed exclusively in a specific neuronal population e.g. dopaminergic neurons or parva-albuminergic neurons in transgenic animals (figure 3b). In 1 mm brain slices with genetically expressed GFP, when cleared by passive CLARITY, tissue transparency reaches a plateau after 5 days, while the fraction of GFP remaining in the tissue decreases rapidly during clearing (Magliaro *et al.*, 2016), demonstrating a trade-off between transparency and protein retention. Antigens and fluorescent proteins are lost during tissue clearing. The optimal clearing is the shortest duration with the best ratio of transparency and protein retention. Magliaro et al. (2016) measured transparency with a regular digital camera and they measured antigen and fluorescent protein loss into the clearing solution with a fluorescent plate reader or a spectrophotometer.

**Figure 3:**
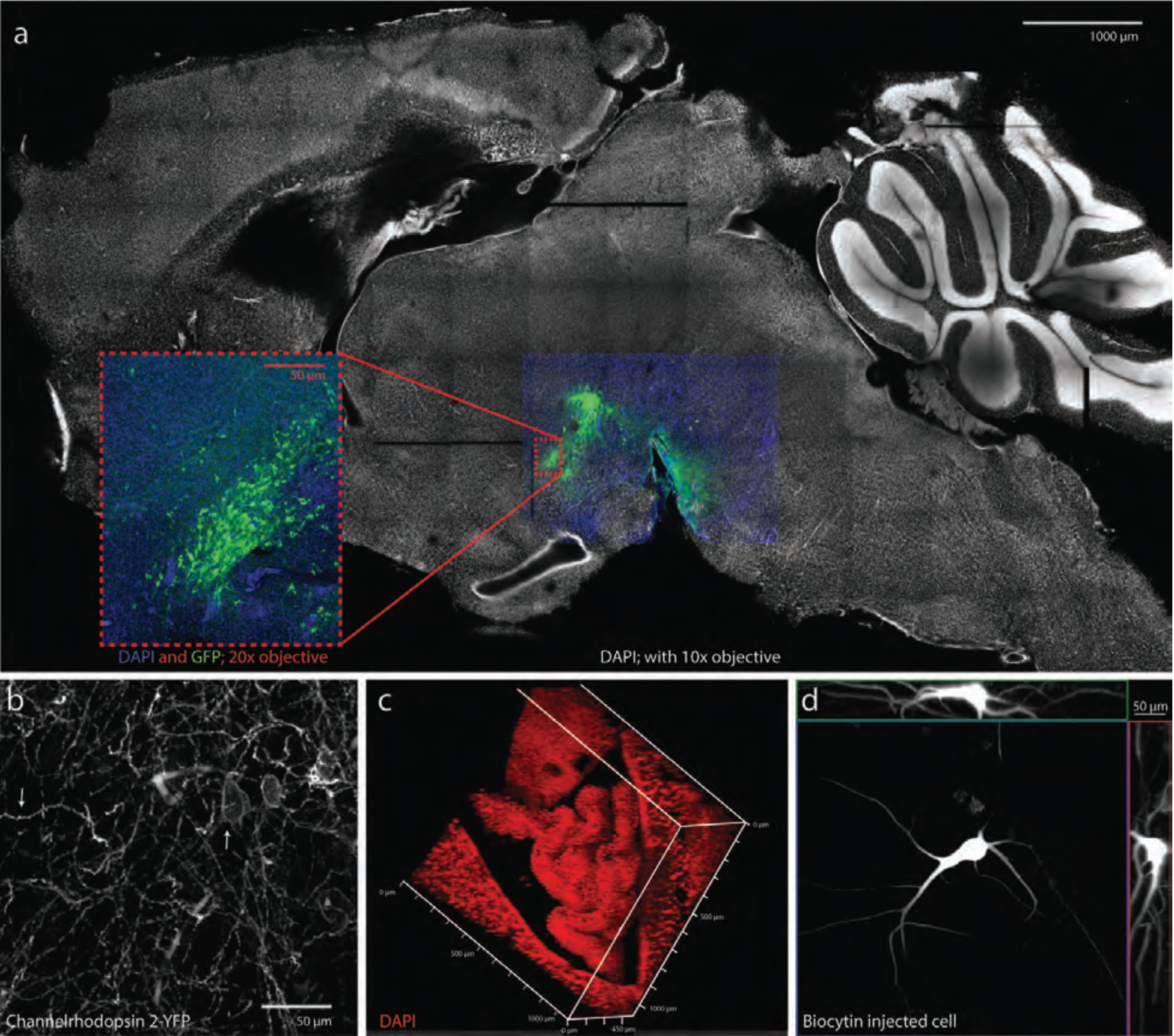
Applications of CLARITY. CLARITY can be used in various purposes. a) Whole field imaging combined with deep tissue imaging (inset) using low magnification (10×) air objects and image stitching. Cell nuclei stained (DAPI: white/blue)of a mid–sagittal mouse brain slice (thickness 2 mm). Low resolution imaging was used to achieve an quick overview of a slab of CLARITY tissue (∼1 min). Viral injected neurons expressing ChR2-YFP in ventral tegmental area was visualised (green). YFP amplified by IHC with a YFP targeting GFP conjugated antibody) (Navntoft, 2015). b) Endogenous expression of yellow fluorescent protein conjugated with channel-rhodopsin-2 (ChR2-YFP) in PV–cells of a CLARITY–treated cortical slice from a transgenic mouse are visible without need of IHC. The neurites (downward arrow) and membranes of the soma (upward arrow). Objective: standard 20× air. Depth ∼300 µm at the maximum of the working distance of 0.55 mm in the 1 mm thick slice. c) The lateral ventricle choroid plexus of mouse reconstructed in 3D after CLARITY and DAPI staining. d) Single neuron staining with biocytin following electrophysiology. Biocytin was injected intracellularly during recording from a motoneuron of the adult turtle spinal cord (Petersen *et al.*, 2014). The neuron was located in a 300 µm slice and 3D–reconstructed with a maximum intensity projection of *Z*(blue), *X*(red) and *Y* (green) plans of the 65 confocal images (*Z*=66 µm) to reveal the morphology of the soma and dendrites. a-d) Tissues were PACT–cleared and imaged in RIMS using a confocal microscope (Zeiss, LSM 700 or 710) with standard 10× or 20× air objectives (Jensen and Berg, 2016).

**Table 2:**
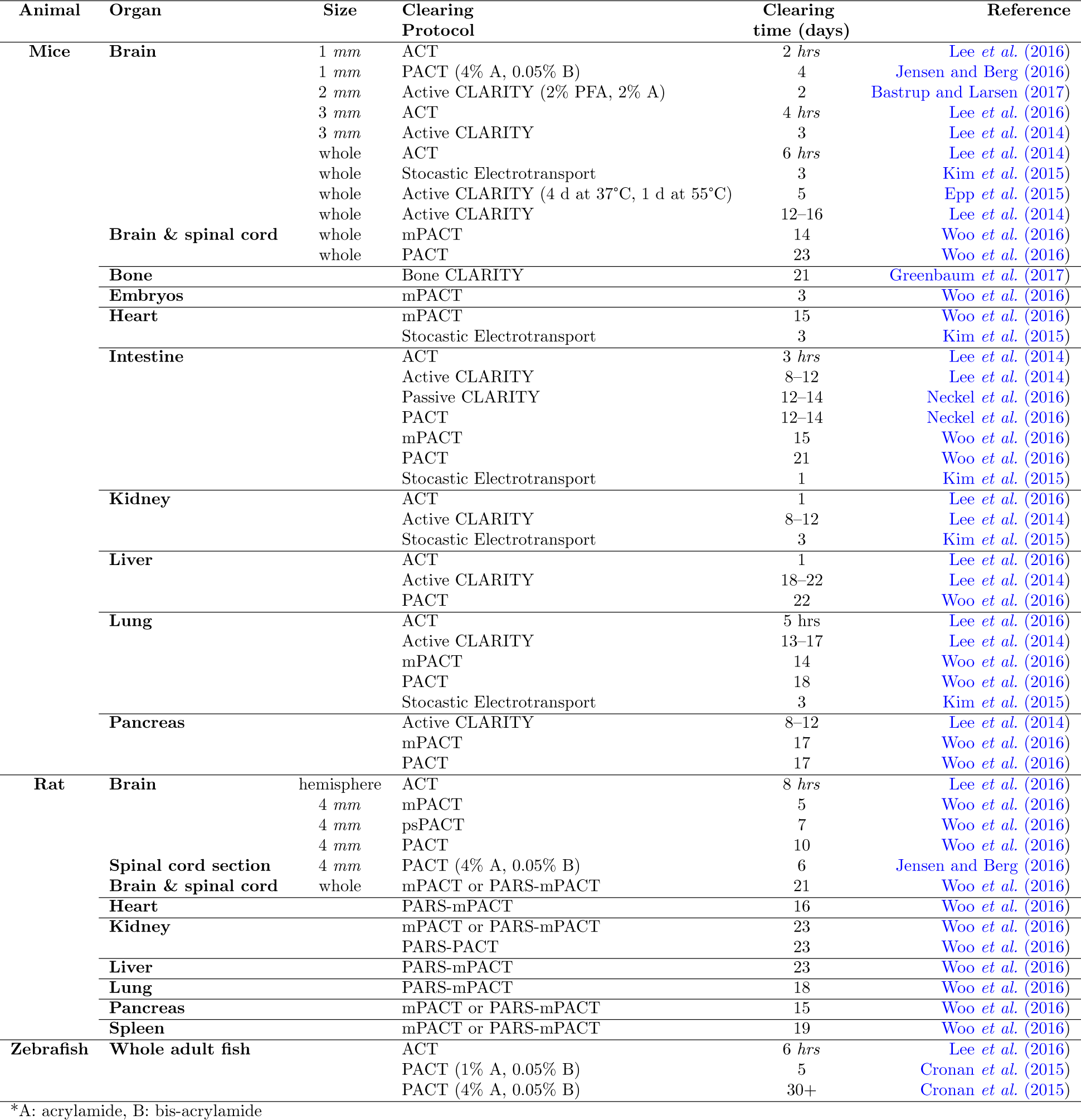
Clearing times for organs and tissues using various protocols. Note that the clearing time do not have a single definition across studies, but is an indication the time to reach sufficient transparency. Organs are whole unless otherwise noted.

## 4. Staining

The main feature of CLARITY is the ability to combine large volume tissue clearing with molecular phenotyping by IHC and other staining methods. Multiple rounds of staining are possible without damage to the preserved structure since the innate biomolecules are chemically bound in the tissue-hydrogel. The tissue clearing solution containing detergent (SDS) can be used to denature antibodies, disrupt binding, and wash antibodies and other molecular labels out of the hydrogel-embedded tissue as preparation for addition rounds of staining. The procedure is simply: incubate overnight with shaking in PBST at 20-40°C, then incubate overnight with shaking in clearing solution at 60°C to wash out antibodies, finally incubate overnight with shaking in PBST at 20-40°C to wash out SDS before next round of IHC and imaging (Tomer *et al.*, 2014). However, there were several shortcomings in this protocol e.g. slow staining, incompatible with lipid dyes and poor retention of RNA, which have since been improved.

### Passive versus active staining

Similar to the clearing of lipids, the application of antibodies for IHC can be performed either passively by diffusion or actively with the assistance of an electrical field or other means. Two of the main challenges in staining are proper uniformity and depth and the penetration of antibodies is relying on diffusion. The slow rate of penetration within the hydrogel is a time–limiting factor in processing samples and thick tissues require weeks to months to adequately label. Increasing the pore size of the hydrogel can be accomplished by reducing acrylamide concentration, but this has limited effect on diffusion speed. The speed is dependent on the molecular weight of the dye or antibody. When choosing a small dye, e.g. DAPI (0.28 kDa), a thick CLARITY-treated tissue can be stained reliably overnight (figure 3a–c), (Jensen and Berg, 2016), whereas staining with heavier antibodies (150 kDa) can take days to weeks. The time of penetration using diffusion can be reduced by half by using smaller antibody fragments such as F(ab’)2 (110 kDa) and Fab (50kDa)) (Li *et al.*, 2015). Another approach to improve antibody staining is using high antibody concentrations or two primary antibodies for a single target, which improved staining speed and quality of parvalbumin-expressing neurons (Bastrup and Larsen, 2017).

The PARS method to clear the tissue can also be used as an active staining method where the staining solution is delivered via the vasculature route mimicking blood flow to reach the whole body and all areas of the tissues to increase speed and penetration depth. If only staining of the brain or CNS is desired delivery by a subdural cannulation directly above the olfactory bulb or in the cisterna magna is more effective.

An initial report demonstrated how an electrophoretically-driven approach could decrease the delivery time of antibodies by taking advantage of their net charge. A static and unidirectional 25 *V* electric field increased antibody penetration by more than 800-fold compared to simple diffusion (Li *et al.*, 2015). Nevertheless, Kim *et al.* (2015) found that static electrophoresis resulted in substantial tissue damage, similar to the problems with ETC when clearing lipids (see above). An alternative strategy was proposed by the Chung group, which is based on *stochastic electrotransport*. The method briefly appeared under the name eTANGO (Richardson and Lichtman, 2015; Hubbert *et al.*, 2014). A rotational electric field selectively dispersed charged molecules without interfering with the endogenous biomolecules. Fluorescent dyes, proteins, and antibodies of different sizes (70-2,000 kDa) was tested using stochastic transport and all gave rapid homogeneous staining of whole brains within a day (Kim *et al.*, 2015).

The benefit of the stochastic electrical field is the more rapid penetration, since speed scales quadratically with the current, and therefore requires a smaller current for the same speed. Hence, the tissue damage is likely to be smaller than for static fields. A static electrical field also has rapid penetration, where the speed scales linearly with the current, but this requires a slightly stronger field for same speed and is therefore more likely to cause damage to the tissue. The main challenge in stochastic electrotransport is building the chamber and buffer flow system. The design is available (www.chunglabresources.com) and a commercial option is expected soon from LifeCanvas Technologies (www.lifecanvastech.com). Lee *et al. (2016)* demonstrated a simpler and cheaper method based on applying pressure to increase the speed and depth of dyes and antibodies penetration into the tissue, which they coined PRESTO (Pressure-Related Efficient and Stable Transfer of macromolecules into Organs) as a part of the ACT-PRESTO protocol. Centrifugal force (c-PRESTO) or convection flow (*s*-PRESTO; for *s*yringe) enabled rapid IHC in 100 µm thick sections within 2–3 h, which compared to passive diffusion requires 1-2 days. *c*-PRESTO requires a standard table-top centrifuge, while *s*-PRESTO requires a syringe pump.

### RNA studies

Strands of RNA are fixed by PFA and retained in the hydrogel, but the potential histological value from RNA is so far unexplored in most clearing methods. Single-molecule fluorescence in-situ hybridization (sm-FISH) of RNA has been demonstrated in thin (100 µm) PACT-processed sections (Yang *et al.*, 2014). By adding a polymerisation step, hybridization chain reaction (smHCR), the depth of RNA detection is extended to 500 µm(Shah *et al.*, 2016), which can visualise e.g. bacterial infections (DePas *et al.*, 2016). However, a modified protocol, EDC-CLARITY, uses carbodiimide chemistry to crosslink the 5’ terminal phosphates to adjacent amines and preserve small RNAs before clearing (Sylwestrak *et al.*, 2016). It uses a less rigid hydrogel (4% PFA, 1% Acrylamide and 0.00625% Bis-acrylamide) and an EDC (1-Ethyl-3-(3dimethylaminopropyl)carbodiimide) fixation step before clearing to retain even microRNA. Depending on the length of the oligonucleotide probe, the probe can be amplified by several strategies i.e. Digoxigenin-labeled locked nucleic acids and tyramide signal amplification that enables measurement of activity-dependent transcriptional signatures, cell-identity markers, and diverse non-coding RNAs in large tissue volumes (Sylwestrak *et al.*, 2016).

### Lipid and membrane stains

Fluorescent dyes, such as the generic *DiI*, *DiD* and other carbocyanine dyes, are lipophilic and therefore primarily stain cellular membranes (Lai *et al.*, 2017). They are extensively used for retro-and anterograde neuronal labelling as well as for marking the position of extracellular electrodes after electrophysiology (Petersen and Berg, 2016) as an alternative to electrical lesioning (Berg *et al.*, 2009). CLARITY and other clearing techniques, such Sca/e and CUBIC (Hama *et al.*, 2011; Susaki *et al.*, 2014) essentially work by washing away the lipids with detergent and solvents. As an unintended consequence, the clearing process therefore also washes out lipophilic dyes, since they adhere to the lipids (Chung *et al.*, 2013; Richardson and Lichtman, 2015; Tainaka *et al.*, 2016). Nevertheless, there are CLARITY-compatible lipophilic and membrane dyes, which can circumvent this problem. DiI—analogues, sulfonated DiI–variants (SP-DiI), and DiI with a chloromethyl benzamide modification (CM-DiI) are aldehyde-fixable to proteins and reliable remain fluorescent in PACT treated tissue (Jensen and Berg, 2016). An alternative is to stain the membranes post-mortem or after tissue-clearing, even with fixable (SP-DiI) or regular DiI (Jensen and Berg, 2016; Xavier *et al.*, 2017). Similarly, the smaller lipophilic FM dyes, are used to image synaptic vesicle exocytosis and endocytosis and has an analogous chemical structure to DiI. FM 1–43FX is a modified FM dye with an aldehyde–fixable aliphatic amine terminal, that also reliably remains and fluoresces stable in PACT treated tissue (Jensen and Berg, 2016).

Since the DiI–analogues are covalently bound to primary amines on proteins by methylene bridges following aldehyde fixation, they are not removed during lipid extraction by solvents or detergents. Other membrane probes that have a aldehyde-fixable anchor point such as mCLING (Revelo *et al.*, 2014) are likely also CLARITY-compatible. Despite the lipid removal and disruption of the lipid membranes, it is possible to perform immunocytochemistry on membrane associated proteins e.g. tight junctions proteins (e.g. Zonula Occludens-1) and channels (e.g. Aquaporin-4) (Neckel *et al.*, 2016).

### Other dyes

Intracellular neuronal labelling is often performed using the generic amide dyes neurobiotin or biocytin. The delivery of such dyes into the neuronal cytoplasm is accomplished either by an intracellular electrode after electrophysiological recording (Petersen *et al.*, 2014; Vestergaard and Berg, 2015; Petersen and Berg, 2016) or by uptake from the nearby surroundings left by juxtacellular deposits in association with extracellular recordings (Wilson and Sachdev, 2004). The dyes are amides and therefore aldehyde-fixable and compatible with CLARITY. Biocytin is not washed out of the cell during CLARITY and can be stained with streptavidin conjugated dye, e.g. Cyanine-3, akin to regular IHC (figure 3d).

CLARITY can also be combined with classical histology such as hematoxylin-eosin and Heidenhain’s azan stain, suggesting potential use in histopathology (Neckel *et al.*, 2016) or combined with colorimetric (nonfluorescent) methods such as horseradish peroxidase conversion of diaminobenzidine to a coloured insoluble product (Sung *et al.*, 2016).

### Minimizing autofluorescence

Two primary sources of autofluorescence in CLARITY treated tissue are heme and lipofuscin. It is therefore important to remove as much blood as possible to reduce the autofluorescent signal from heme. Under normal conditions, blood is removed during the initial cardiac perfusion, but in special situations where perfusion is not possible, such as in human tissue, heme can be eluted by incubating hydrogel-embedded PACT sections in aminoalcohol (CUBIC reagent-1: mixture of 25% (w/v) urea, 25% (w/v) N,N,N’,N’-tetrakis(2-hydroxypropyl) ethylenediamine and 15% (w/v) Triton X-100 (Susaki *et al.*, 2014), or 25% (w/v) N,N,N’,N’-tetrakis(2-hydroxypropyl) ethylenediamine in PBS alone for 12-24 h at 37°C while shaking before (Treweek *et al.*, 2015) or after clearing (Greenbaum *et al.*, 2017). Lipofuscin autofluorescence is partially countered by the tissue clearing process. However, when the tissue sections are thick, they may be incubated in 0.2%-1.0% (w/v) Sudan Black B, which is a nonfluorescent lipophilic dye, in 70% ethanol for 1-3 hours immediately before hydrogel-polymerisation in order to further reduce autofluorescence (Treweek *et al.*, 2015).

## 5. Optical Clearing

The final step before imaging is optical clearing, where the average refractive index (RI) of the hydrogel (∼1.46) and the imaging solution are closely matched. Photons from both the excitation light and the emitted fluorescence signal scatter if the RI is not matched when travelling through the sample, which limits the quality and depth of imaging. The RI of each solution can be matched to the optimal RI of the tissue or microscope objective by adjusting the concentration of the main ingredient, e.g. Histodenz. Brain tissue usually has an average RI of 1.46-1.47, whereas e.g. bone has 1.48-1.49. It is possible to image without a mounting solution, but in simple water or PBST e.g. PBST (0.1% Triton X-100 in PBS) (Poguzhelskaya *et al.*, 2014). However, the tissue remains swollen, and water and PBST have a lower RI (∼1.33) than of the hydrogel (∼1.46) and therefore the tissue sample will appear opaque or cloudy. Imaging in water is part of the ePACT method where the tissue is intentionally expanded for greater resolution. There are several different options of mounting solutions (table 4).

### FocusClear: the generic imaging solution

The original FocusClear (CelExplorer Labs Co.) with the main ingredient diatrizoic acid has an RI of 1.454 similar to the tissue-hydrogel, but is expensive (∼29 USD/ml), and not usable for storage (Tomer *et al.*, 2014). If the sample is not washed properly in PBST, a *irreversible* white precipitate, which is likely caused by a reaction with remaining SDS, can develop within the embedded tissue after a few days.

**Table 3:**
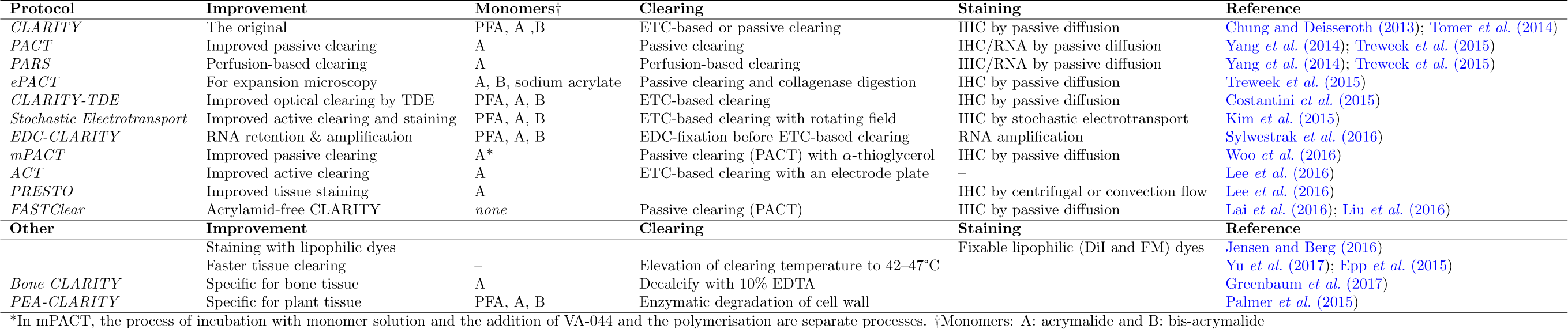
Different protocols and improvements based on CLARITY

**Table 4:**
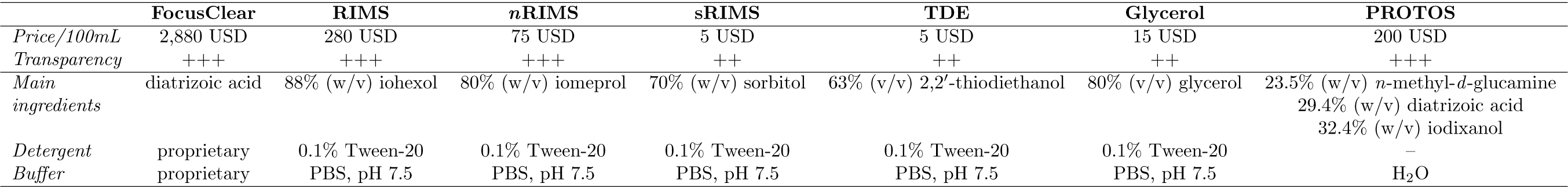
Imaging solutions. Solutions for adjusting refractive indices and their transparency, cost, main ingredients, detergents and buffers. Prices are approximate.

### RIMS and nRIMS: the optimal solutions

Yang *et al.* (2014) developed an affordable alternative to FocusClear coined *r*efractive *i*ndex *m*atching *s*olution (RIMS) of 88% (w/v) Histodenz in PBS with an RI of 1.46. Both Diatrizoic acid and Iohexol (HistoDenz) are complex molecules containing an aromatic ring and three iodine atoms, which provides a large number of electrons for interaction with passing light waves, i.e. high refractive index, but in a relatively low-concentration and low-viscosity solution (Richardson and Lichtman, 2015). Woo *et al.* (2016) used a 80% (w/v) solution of Iomeprol-d3 (also known as Nycodenz and Iohexol-d5), referred to as *n* RIMS, which also has an RI of 1.46. A similar clearing solution is PROTOS (Murray *et al.*, 2015; Kim *et al.*, 2015). RIMS reportedly works better for rodent brains and is reportedly less ideal for human brain tissue, where 47% TDE in PB is better (Liu *et al.*, 2015).

### Glycerol, Sorbitol and TDE: affordable alternatives

Glycerol, sorbitol, and thiodiglycol (2,2’-thiodiethanol, TDE) are water-soluble and low-viscosity liquids that can be used to tune an aqueous solution over a range of refractive indices by dilution in water. Glycerol and sorbitol are inexpensive and are often found in conventional clearing reagents and in mounting media. TDE was initially used in mounting media for super-resolution microscopy (Staudt *et al.*, 2006), but concentrations of 40-60% (v/v) can also clear large tissues i.e. rodent brains (Costantini *et al.*, 2015; Aoyagi *et al.*, 2015). Solutions with 63% (v/v) TDE, 80% (v/v) glycerol, or 70% (w/v) sorbitol (coined sRIMS) all have an RI of ∼1.46. One drawback to TDE is that at high concentrations it reduces the brightness of some green fluorophores (Staudt *et al.*, 2006). TDE (47%) in PB (instead of PBS) is reportedly better for RI-matching in human tissues (Liu *et al.*, 2015). RIMS outperformed sRIMS and glycerol regarding imaging resolution and depth (Yang *et al.*, 2014; Marx, 2014). Another option is a mix of DMSO and *D*-sorbitol (Economo et al., 2016).

### Storage of CLARITY tissue

All solutions apart from FocusClear can be used for storage. However, a slight loss of fluorescent signal of AlexaFluor-568 has been reported in glycerol (Liu *et al.*, 2015), although the fluorescent signal loss of (red) DiI-dyes has not been observed over 6 months in glycerol (Jensen and Berg, 2016). Storage at room temperature as opposed to using refrigeration is recommended since precipitate can appear at lower temperatures. All solutions should include 0.01% sodium azide to prevent bacterial and fungal growth. Lowering the PBS concentration to 0.005 M phosphate buffer reduces the appearance of salt precipitate at colder temperatures i.e. 4°C(Treweek *et al.*, 2015).

## 6. Imaging CLARITY tissue

Imaging of the clarified tissue is more difficult than conventional histological samples. The point of the lipid removal is to make the tissue transparent for the purpose of imaging in depth. This changes the task from imaging single or several 2-dimensional (2D) sections to a 3-dimensional (3D) volume, which is a much larger volume than what is used in conventional microscopy (figure 3c-d). Such imaging requires considerations of imaging time and resolution when selecting the microscope and objective as well as sample size.

### Choice of microscope

Standard light microscopy is generally not suitable for large transparent tissues samples as the excitation light penetrates the sample and generates fluorescence signal from the whole sample. The signal from the focused plane will be attenuated and likely lost by the fluorescence coming from elsewhere. While it is possible to image highly fluorescent neurons (600 µm deep in 2 mm tissue) such image will require postprocessing deconvolution (Epp *et al.*, 2015). However, a low magnification (i.e. 1.5-5×) wide field or stereo microscope can be helpful in an initial examination of samples for identifying fluorescence signals from e.g. labelled electrode traces before further investigation (Jensen and Berg, 2016; Petersen and Berg, 2016).

Imaging of CLARITY-processed tissue samples can be performed using standard confocal microscopes. No modifications are required, but optimal choice of objectives is recommended (see below). However, processing time is slow, especially for high resolution, high sampling, and large volumes, furthermore, illuminates of the entire sample during imaging can lead to photobleaching during long imaging sessions and when imaging at large depths.

The properties of two-photon microscopy lead to significant lower photobleaching of fluorophores and may also provide greater imaging depths than confocal microscopy. The serial point–by–point scanning in 3D tissue is similarly slow.

Light-sheet microscopy is a 3D imaging technique is ideal for clarified samples. Light-sheet microscopy achieves optical sectioning by selectively confining the illumination to the plane of interest by using a Gaussian or a Bessel beam from the side of the tissue. Furthermore, while confocal and two-photon microscopy is point scanning, and hence inherently slow, light-sheet microscopy uses fast sCMOS or CCD camera sensors to image the selectively illuminated focal plane, resulting in minimal photobleaching and increased imaging speeds that are 2–3 orders of magnitude faster than confocal and two-photon microscopy (Tomer *et al.*, 2014).

### Microscope objectives

Objectives with high numerical aperture and long working distances are desirable for maximum imaging depth and resolution. Furthermore, objectives should be optimised for an RI of 1.46 for CLARITY cleared tissues. Several objectives have been developed specifically for CLARITY (table 5).

When using a non-optimized objective, it is best to use the one designed for the RI near 1.46 e.g. glycerine (RI 1.47) or oil immersion (RI 1.52). Water-immersion lenses (RI 1.33) work better than air objectives (RI 1). Furthermore, it is time-consuming imaging a large area using a 25× objective, since this requires capturing and stitching of multiple tiles of images. Such strategy also increases the risk of photobleaching. A lower magnification 10x can be more useful to image over larger areas (figure 3a), and they have longer working distances. Indeed, the affordable and commonly available low magnification (10-20×) air lenses are compatible and have been successfully used for several CLARITY studies (Jensen and Berg, 2016; Hsiang *et al.*, 2014; Poguzhelskaya *et al.*, 2014; Bastrup and Larsen, 2017). Images in figure 3 were captured with low magnification (10-20×) air lenses. CLARITY-based tissue clearing provided an increased signal-to-noise ratio, and staining homogeneity in super-resolution stimulated emission depletion (STED) microscopy (i.e. with a 100× objective) of kidney tissue (Unnersjö-Jess *et al.*, 2015). During extended imaging moisture can evaporate from the imaging medium and cause subtle changes RI and resulting in aberrations and loss of resolution. A hygroscopic imaging media or sealed chamber can prevent this.

**Table 5:**
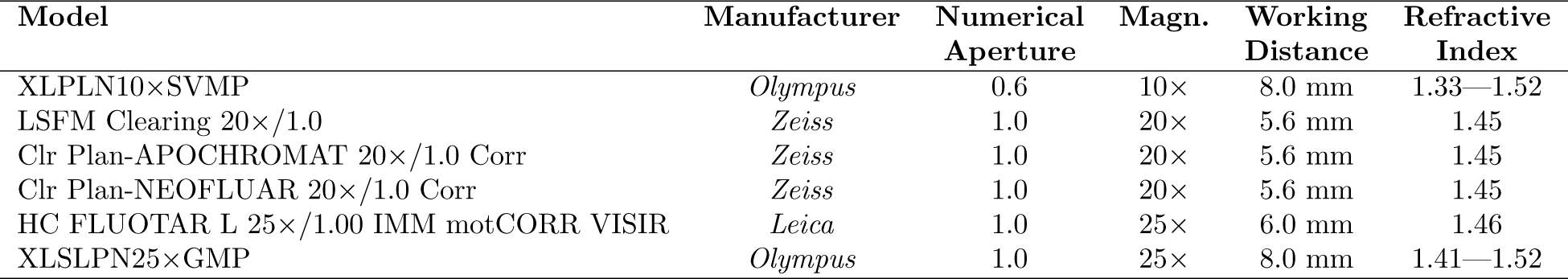
Microscope objectives optimised for imaging CLARITY samples

### Stereology and tractography

Quantification of total cell numbers, densities and projections in tissue while preserving spatial information has been a challenge in *stereology*, since it relies on interpretation of 2D sections of tissues and statistical sampling methods from several histological tissue sections (Gundersen and Jensen, 1987; Walloe *et al.*, 2011).

Applying stereological methods to cleared tissues eliminates the need for labour-intensive sectioning. Furthermore, clearing and counting in the whole 3D volumes rather than multiple sampled 2D sections allows for better stereological estimates. Stereology in CLARITY tissue also has the advantage of the ability to detect subtle changes that might be overlooked because of sampling variance (Erskine and Khundakar, 2016; Greenbaum *et al.*, 2017; Bastrup and Larsen, 2017). However, the computational challenge of aligning images, down-sampling, and creating a 3D–visualisation are currently only semi-automated and for many investigators such task represents a strain. The performance of automated cell detection and segmentation algorithms as alternatives to manual stereological cell counting are still limited by lower detection rates and higher false-positive rates (Schmitz *et al.*, 2014).

Similarly, quantifying axon tracts is also a challenge. Therefore, Deisseroth and colleagues developed a method to compute 3D structure tensors from CLARITY images using tools adapted from diffusion–MRI tractography (Ye *et al.*, 2016). Ye *et al.* (2016) used activity dependent viral expression of fluorescent proteins to quantify axonal projections of behaviourally defined neuronal populations. They termed this method *C*LARITY-based *a*ctivity projection tracking *u*pon *r*ecombination (CAPTURE). The method requires viral tools and light sheet imaging, but software for this large–scale image analysis is freely available at captureclarity.org.

### Author recommendations

Several qualities are worth considering when selecting or optimising a CLARITY protocol (figure 4). From our experience we recommend the following:

**Figure 4:**
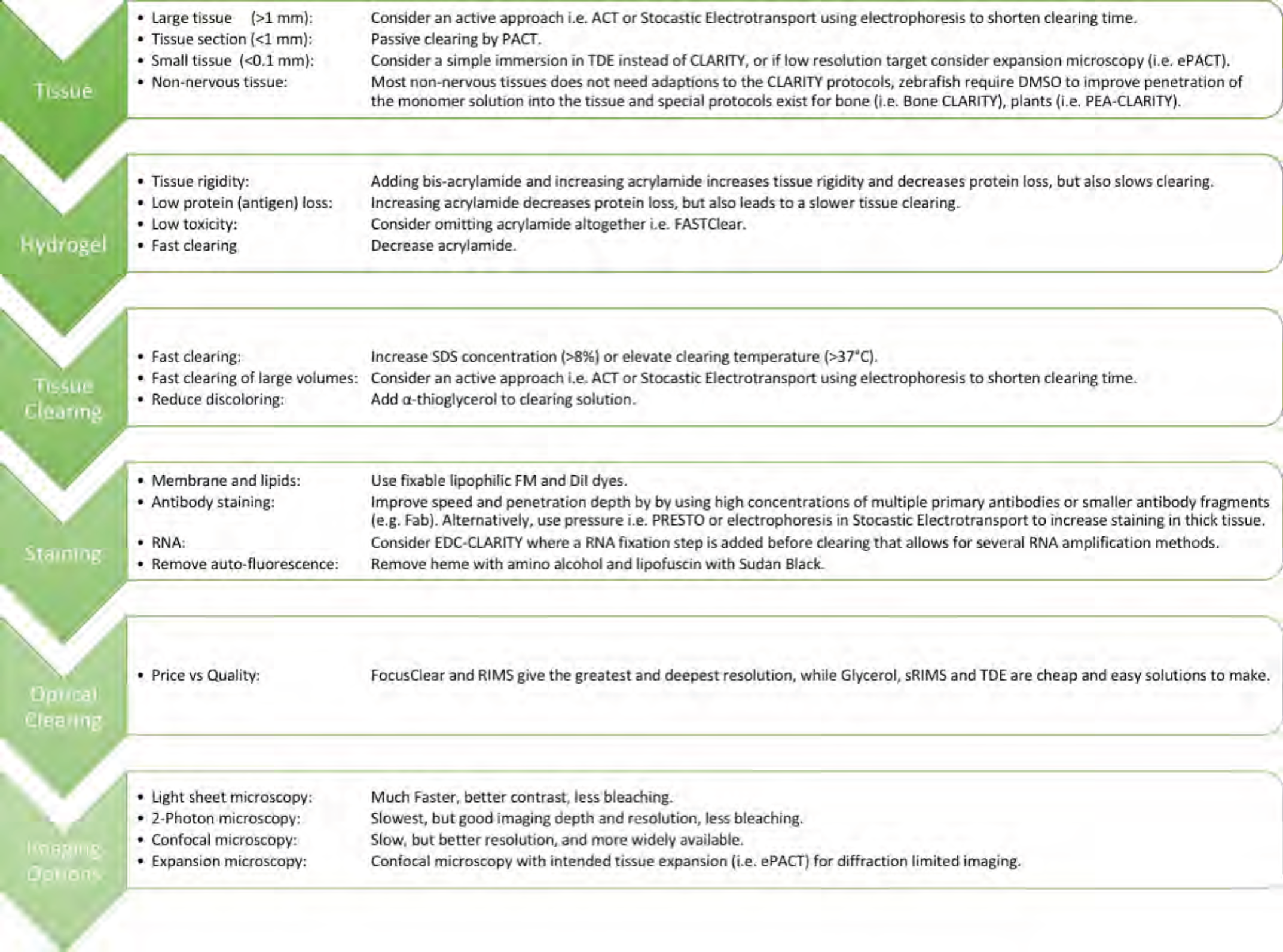
Factors and qualities when selecting and optimising a CLARITY-protocol.

**For most tissue use PACT or mPACT:** The passive protocols are uncomplicated and inexpensive to setup for novel users. The protocols that omits PFA result in faster clearing with less protein loss and better staining. However, we also recommend including 0.05% bis-acrylamide in the monomer solution because it makes for a more rigid hydrogel. The solidification of the monomer solution surrounding the tissue is an indicator of successful polymerisation, although bis-acrylamide may slow the clearing. Clearing speed can further be improved by elevating the temperature during clearing to 42-47°C and employing mPACT. A minimalist incubation chamber can consist of a styrofoam box with a temperature–controlled heating pad, and then placing the box and a gentle lab shaker.

**For large tissues use ETC:** For active clearing of large tissues with a simple ETC chamber with single wires for thick tissue sections (1-3 mm), we recommend the design by Bastrup and Larsen (2017) where the distance between wires can easily be adjusted and enables tissue clearing of variable tissue sizes while minimising the resistance and limiting the heat generated in the system. For whole brains or organs, we recommend the more elaborate stochastic electrotransport approach. The addition of 0.5% *α*-thioglycerol to prevent tissue discolouration during electrophoresis recommended regardless of chamber design.

**Use glycerol and RIMS:** Selecting an imaging solution is a matter of price versus quality. We regularly use 80% glycerol and RIMS for imaging and storage. See table 4 for options of mounting solutions.

## Successful applications using CLARITY

CLARITY is applicable not only to nervous tissue, but in principle to any biological tissue. The spinal cord is enveloped by dense grey matter, and the myelin can be difficult, although not impossible, to clear, and limits access to the internal white matter (Jensen and Berg, 2016; Spence *et al.*, 2014; Liang *et al.*, 2015, 2016). One simple solution is to split the spinal cord down the midline into two hemisections, which may be easier to stain and image than using PARS-CSF. While CLARITY was originally demonstrated in mice and rat brains, CLARITY and its variants have been used in several other tissues and organisms. Zebrafish is a popular neuroscience model organism and has been cleared in PACT, but with an added 5% DMSO to the monomer solution to increase penetration as well as 1% PFA and 0.05-0.025% bis-acrylamide to increase rigidity (Cronan *et al.*, 2015; Frétaud *et al.*, 2017).

### Human samples

Application of CLARITY on human brain samples opens the possibility of visualising human pathology in a novel way. Transcardial perfusion is obviously impossible and clinical samples are usually fixed and stored in formalin over extended periods. Indeed, the human pathology of Alzheimer’s (Ando *et al.*, 2014), Parkinson’s (Liu *et al.*, 2015), neurodegeneration due to mitochondrial disease (Phillips *et al.*, 2016), autism (Chung *et al.*, 2013) as well as intractable epilepsy (Costantini *et al.*, 2015) have been visualised in 3D by CLARITY on formalin-fixed tissues. A frozen brain sample, which was PFA-fixed and cryoprotected in sucrose, has also been cleared and stained using CLARITY (Phillips *et al.*, 2016). The human enteric nervous system was also studied recently using CLARITY (Neckel *et al.*, 2016). The speed of tissue clearing differs between CNS regions depending on the degree of myelination and the duration of formalin fixation. The transient tissue expansion during CLARITY-clearing of animal tissue has been observed to be irreversible in human brain tissues, especially after prolonged (>40 days) passive tissue clearing (Liu *et al.*, 2015). Lai *et al.* (2016) tested an acrylamide-free version of CLARITY, i.e. essentially only washing with SDS followed by RIMS, and found it gave better results for formalin-fixed tissue. They argued that the previously suggested tissue-PFA-acrylamide cross-linking does not actually occur. For this reason, omitting acrylamide is better and they named this protocol Free of acrylamide sodium dodecyl sulphate (SDS)-based tissue-clearing (FASTClear) (Liu *et al.*, 2016).

## Beyond the CNS

The enteric nervous system and mesenteric vasculature can be visualised in 3D with CLARITY (Neckel *et al.*, 2016). While it is possible to clear skeletal muscle with CLARITY, labelling the neuromuscular junctions with fluorescently labelled *α*-bungarotoxin has not been possible, likely due to the cross-linking and fixation preventing the access of the toxin to the acetylcholine receptors (Milgroom and Ralston, 2016). However peripheral nerves could be targeted by IHC or an anterograde lipophilic tracer e.g. SP-DiI or a viral tracer instead. It remains to be tested whether labelling of Isolectin B4-positive nociceptor cells is compatible with CLARITY clearing. Neurovasculature has also been studied, where the tissue is incubated in or flushed with antibodies through the vasculature, i.e. anti-CD31 (Neckel *et al.*, 2016; Woo *et al.*, 2016). The endothelial cells could also be stained by flushing the vasculature with a DiI-dye or biocytin before fixation.

### Muscle and bone

It is possible to clear skeletal, cardiac and smooth muscle with CLARITY-based clearing with no modifications to the protocols (Yang *et al.*, 2014; Epp *et al.*, 2015; Milgroom and Ralston, 2016; Gloschat *et al.*, 2016; Kolesová *et al.*, 2016; Sung *et al.*, 2016; Ding *et al.*, 2017). However, the collagen-rich tendons are difficult to make transparent, similar to the myelin-rich white matter of the spinal cord. Extended clearing time and adjustment of the RIMS RI may improve tendon transparency. Chondrocytes can be stained with the fixable lipophilic dye, DiI-SP (Jensen and Berg, 2016; Calve *et al.*, 2015). Bone tissue, however, is challenging since it has low lipid content, a hard mineral component in addition to the soft bone marrow. Clearing osseous tissue can be done using solvent based methods (Richardson and Lichtman, 2015; Greenbaum *et al.*, 2017). Bone tissue can also be cleared using CLARITY by adding a decalcification step, PACT-deCAL, in which tissue is placed in a 0.1 M EDTA (in PBS, pH 8 for 2 days) during PACT-clearing with increased SDS concentration and pH (8-10% SDS and pH 8). Nevertheless, this gave only modest visualisation depth of 200-300 µm(Treweek *et al.*, 2015). By extending and increasing the decalcification step (0.3 M EDTA and 14 days) Greenbaum *et al.* (2017) was able to extend visualisation depth to 1.5 mm in a protocol called ‘Bone CLARITY’. The heme-rich bone marrow also presents a problem regarding autofluorescence, which has be reduced by removing heme with amino alcohol before refractive index matching (figure 5). The osteoblasts can be stained with the fixable lipophilic dye CM-DiI (Jensen and Berg, 2016; van Gastel *et al.*, 2012).

**Figure 5:**
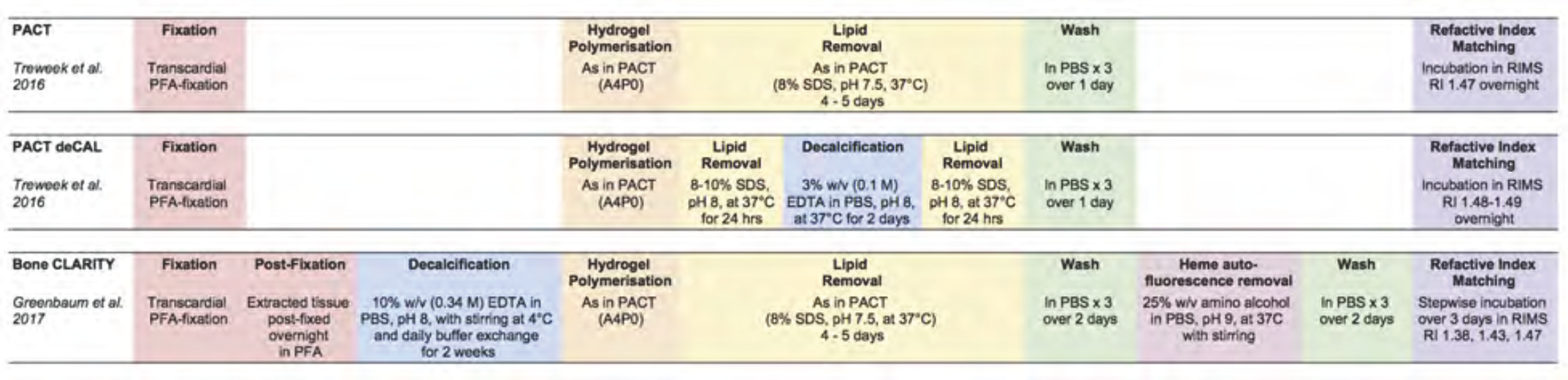
Bone tissue clearing by modified CLARITY protocols. The steps of the three basic protocols for clearing bone tissue are listed in horizontal direction. The essential modification of the original PACT (top) is the addition of a decalcification step (blue) in PACT deCAL (middle) and Bone CLARITY (bottom). The latter also includes reduction of autofluorescence.

### Other organs and species

CLARITY-based protocols have been used with little, e.g. lower acrylamide concentrations, or no modifications to clear most other organs such as lever (Lee *et al.*, 2014; Font-Burgada *et al.*, 2015), kidney (Yang *et al.*, 2014; Lee *et al.*, 2014; Unnersjö-Jess *et al.*, 2015), pancreas (Lee *et al.*, 2014; Muzumdar *et al.*, 2016), adrenal gland (Epp *et al.*, 2015), spleen (Epp *et al.*, 2015; Kieffer *et al.*, 2017), lymphoid tissues (Kieffer *et al.*, 2017), intestine (Yang *et al.*, 2014; Lee *et al.*, 2014; Epp *et al.*, 2015; Neckel *et al.*, 2016; Kieffer *et al.*, 2017), testes (Epp *et al.*, 2015; Frétaud *et al.*, 2017), ovaries (Feng *et al.*, 2017), and lung tissue (Yang *et al.*, 2014; Lee *et al.*, 2014; Epp *et al.*, 2015; Saboor *et al.*, 2016). However, given that organs have different densities and ratios of connective tissue, lipids, and protein the clearing times and the average RI of the tissue-hydrogel vary (table 2). For spleen and kidney, the best RI-matching solutions were found to be CUBIC-mount, RIMS, sRIMS and TDE, respectively (Lee *et al.*, 2016).

Vertebrate and invertebrate animal models, such as rabbits, chickens, zebrafish, xenopus, small octopuses were cleared by the ACT (Lee *et al.*, 2016). Other models include sea lamprey (Chung-Davidson *et al.*, 2014) and turtles (figure 3d). Adding an enzymatic degradation step as in Plant-Enzyme-Assisted (PEA)CLARITY, it is possible to remove the plant cell walls after tissue-clearing in SDS to clear and stain whole plant tissues without the need for any sectioning of the material. This can be useful in localisation of protein in intact plant tissue in 3D while retaining cellular structure (Palmer *et al.*, 2015). CLARITY has also been used with hybridization chain reaction to detect rRNA from and visualise bacterial infections (DePas *et al.*, 2016).

## Alternative clearing methods

There are several new and old alternatives to CLARITY with their own advantages and limitations. Tissue-clearing methods can be divided into three groups: *organic solvent–based*, *aqueous–based* and *hydrogel-based* (Figure 6).

**Figure 6:**
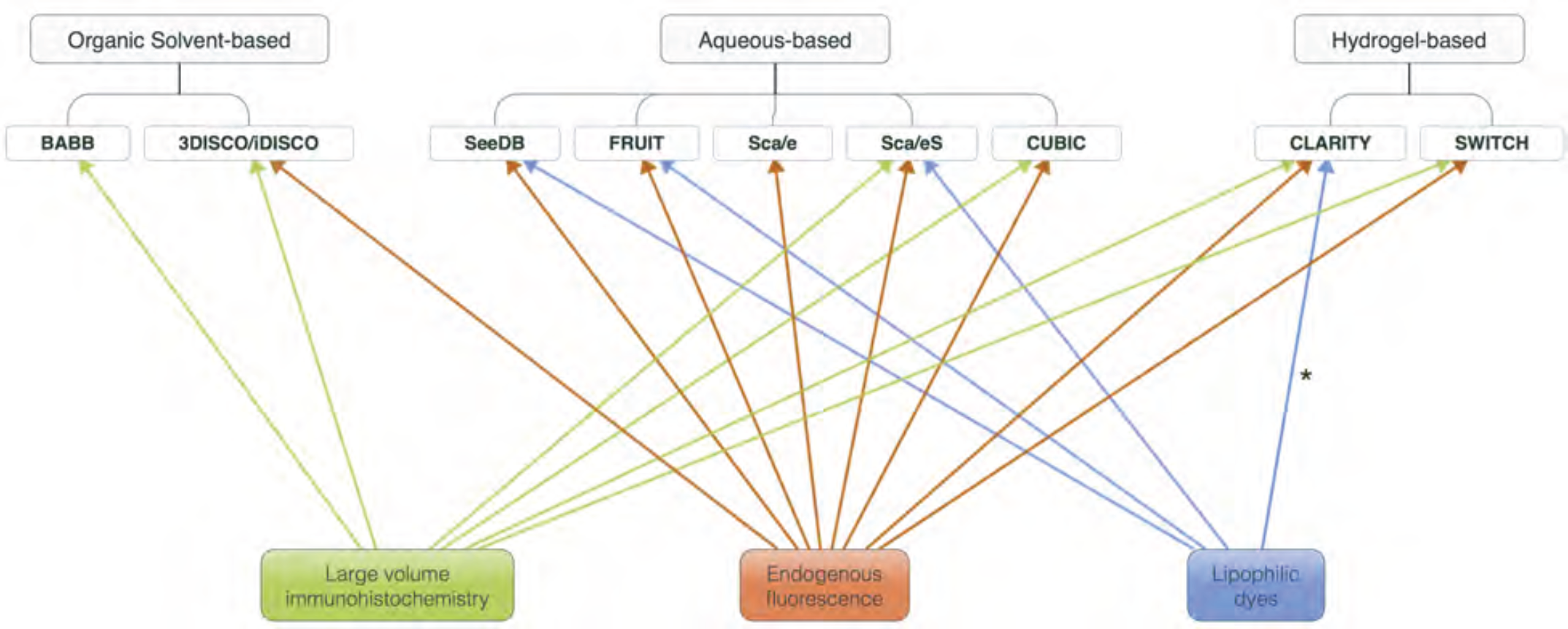
Clearing methods and ideal applications. Clearing methods fall into three categories: organic solvent–based (top, left), aqueous–based (top, middle) and hydrogel–based (top, right). Each method have advantages and shortcomings. The choice of approach should be guided by the histological question. Histological questions are connected with one or more applications listed below: Large volume immunohistochemistry (green), preservation of endogenously expressed fluorescent proteins (red) or tracing using lipophilic dyes (blue). The compatibility of topics with individual methods are indicated by the arrows. For instance, endogenous expression of fluorescent proteins is not ideal to be studied with BABB and 3DISCO, since these methods quenches fluorescence within hours (BABB) or days (3DISCO). *Modified dyes are required since traditional lipophilic dyes are washed out during CLARITY.

### Organic solvent–based

The first tissue clearing method by Spalteholz a century ago was based on the replacing water within the tissue with a mixture of organic solvents i.e. benzyl benzoate and methyl salicylate, which improves transparency by reducing light scattering (Spalteholz, 1914). Since benzyl benzoate is insoluble in water, it required an intermediate dehydration step with ethanol. Subsequent organic solvent–based methods have the same scheme i.e. dehydration with lipid solvation followed by an alcohol/ether–step then followed by additional lipid solvation–step and RI-matching (Richardson and Lichtman, 2015). Dodt *et al.* (2007) refined the method by employing mixture of benzyl alcohol/benzyl benzoate (BABB). However, BABB quenches the emission of fluorescent proteins within hours (Richardson and Lichtman, 2015), and the gentler solvents tetrahydrofuran (THF) and dibenzylether (DBE) in the 3DISCO method prolongs the emission for 1-2 days (Becker *et al.*, 2012; Ertürk *et al.*, 2012). This approach has an advantage in rapid clearing kinetics due to the quick diffusion of small molecules and imaging speed due to a tissue shrinkage. 3DISCO has limited antibody penetration depth (250 µm)(Hirashima and Adachi, 2015), but has been improved in the iDISCO protocol (Renier *et al.*, 2017). Organic solvents are generally toxic and some, i.e. THF and DBE, are even explosive.

### Aqueous-based

Aqueous-based clearing solutions work by detergent-based lipid removal and hydration. The RIs are then matched in an aqueous solution of concentrated sugar– or contrast–agents with a high RI (1.44–1.49). The RI-matching is accomplished by the inclusion of glycerol in Sca/eA and Sca/eS (Hama *et al.*, 2011, 2015) or sugars e.g. fructose or sucrose in high concentration in CUBIC, SeeDB, FRUIT (Susaki *et al.*, 2014; Ke *et al.*, 2013; Hou *et al.*, 2015). Some protocols use urea to enhance the hydration of biomolecules and improve penetration into the tissue i.e. Sca/e, SeeDB, CUBIC (Hama *et al.*, 2011; Susaki *et al.*, 2014; Ke *et al.*, 2013), although this can lead to tissue swelling. They are better at preserving fluorescent proteins compared with organic solvent–based clearing. Generally, aquous–based clearing induces limited change in tissue size, and the compounds are relatively harmless, but the protocol can be labour intensive.

SeeDB and FRUIT provide good transparency, but they have limited permeability for macromolecules and are therefore incompatible with IHC (Ke and Imai, 2014; Hou *et al.*, 2015). The gentler methods omitting strong detergents, glycerol and amino alcohols (i.e. Sca/eS, SeeDB and FRUIT) are however compatible with lipophilic dyes (Hama *et al.*, 2015; Ke *et al.*, 2013; Hou *et al.*, 2015). Yet, fixable lipophilic dyes are likely compatible with all aqueous-and solvent-based methods (Jensen and Berg, 2016).

## Hydrogel-based

Hydrogel-based clearing is innovative approach, that works similar to aqueous-based methods, but has a more aggressive lipid removal combined with impregnation of the tissue with a monomeric component (Silvestri *et al.*, 2016). Polymers and gels, e.g. paran and OCT compound, have long been used to support tissues during sectioning and histology. Hydrogels have been a focus of tissue engineering as a substrate for cell culture and a scaffold for growing organs (El-Sherbiny and Yacoub, 2013). In contrast, in the hydrogel-based clearing, the supporting gel is built from within the tissue by anchoring the cellular components to the gel monomers. Cross–linking produces a polymerization, where the unlinked elements, such as lipids, can be removed and give room for macromolecular probes and IHC (Deisseroth, 2017; Silvestri *et al.*, 2016). Since 2013, many variations and improvements of the original hydrogel-based clearing concept, CLARITY, have emerged, which we reviewed above. However, two divergent approaches are briefly mentioned here.

## SWITCH

Using a *s*ystem-*w*ide control of *i*nteraction *t*ime and kinetics of *ch*emicals (SWITCH) Murray *et al.* (2015) replaced the original polyacrylamide mesh with a more stable dialdehyde, i.e. glutaraldehyde. Glutaraldehyde can penetrate through large tissues at pH 3, circumventing the need for tissue perfusion, and rapidly initiate cross-link reactions deep inside the tissue when the pH is *switched* to pH 7. After fixation, lipids are removed by boiling the tissue at 60–80°Cin abuffer with SDS. The rigid fixation also preserves some lipids and small molecules such as dopamine after clearing, and the harsh treatment does not result in significant loss of proteins or antigens. A similar on/off control of the binding kinetics of the antibodies improves the penetration antibodies. SWITCH also allows for iterative IHC staining as CLARITY, but it is more stable and allows up to 20 rounds of IHC.

### Expansion Microscopy

This technique transforms small tissue samples or cell cultures into swellable hydrogels similar to that of CLARITY by adding the super–absorbent sodium acrylate copolymer (Chen *et al.*, 2015). The purpose is to study small species without the need for tissue–clearing since the sample is expanded and therefore transparent. The hydrogel serves as a scaffold that is partially enzymatically digested and expanded by almost two orders of magnitude to improve resolution beyond the optical diffraction limit.

## Conclusion

Much work has increased the clearing speed of the method CLARITY and adapted its use to several tissue types i.e. non-nervous tissue, bones, and plants. This increases our understanding of biology in different organs and species. While IHC in large volumes continues to be difficult, the main limitation seems to be in imaging and data processing of the large volumes. Light-sheet microscopy makes image–acquisition fast and easy, but is not readily available to all researchers. Furthermore, large image volumes give large data volumes (giga-to terabyte) that are difficult to handle and analyse. The challenge of image acquisition and analysis, therefore, lies beyond CLARITY and is a problem shared by all tissue clearing methods. Improvements in large volume image data analysis will improve not only help experimenters using CLARITY, but bring histology into the third dimension.

## Acknowledgements

We thank Editor-in-chief Dr H. Steinbusch for the invitation for this review. Charlotte A. Navntoft for injection of viral tracers, Robertas Guzulaitis for biocytin injection, and Rasmus Christensen and Tania Barkat for providing rodents. Thanks to the Core Facility for Integrated Microscopy, Faculty of Health and Medical Sciences, University of Copenhagen, and the assistance provided by Thomas Braunstein and Laura Plantard. The work is part of the Dynamical Systems Interdisciplinary Network, University of Copenhagen, and supported by the Danish Council for Independent Research medical science (R.W.B).

